# Chlamydial membrane vesicles deliver the chlamydial porin OmpA to mitochondria to inhibit apoptosis

**DOI:** 10.1101/2025.06.02.657351

**Authors:** Andreea Mesesan, Henry Oehler, Collins Waguia Kontchou, Aladin Haimovici, Martin Helmstädter, Oliver Kretz, Oliver Schilling, Irina Nazarenko, Ulf Matti, Jonas Ries, Ian E. Gentle, Georg Häcker

**Author notes:** These authors contributed equally (Andreea Mesesan, Henry Oehler).

## Abstract

*Chlamydiae* are obligate intracellular bacteria that inhibit mitochondrial apoptosis to maintain integrity of the host cell. We have previously reported that a chlamydial outer membrane β-barrel protein, the porin OmpA, can during ectopic expression inhibit mitochondrial apoptosis through direct interaction with the BCL-2-family effectors BAX and BAK. We here show that OmpA from *Chlamydia trachomatis* (*Ctr*) uses membrane vesicles for its delivery to the outer mitochondrial membrane during *Ctr* infection. Using a number of imaging and fractionation techniques, we show that OmpA during infection reaches mitochondria and is inserted into mitochondrial membranes. Purified membrane vesicles (MV) from *Ctr*-infected cells contained OmpA. When added to uninfected cells, MV fused with mitochondrial membranes, causing the interaction of OmpA with BAK and the cytosolic retro-translocation of BAX. MV addition to uninfected cells protected the cells against apoptosis. We propose a structural model of this BAK inhibition by OmpA that reenacts the inhibition of BAK by the mitochondrial porin VDAC2. The results provide evidence that the porin from *Chlamydia*, as well as the structurally similar porin from the related *Simkania*, specifically exploits its relationship to mitochondrial porins to protect the infected cell against apoptosis and to enable intracellular growth of the bacteria in human cells.

## Introduction

*Chlamydia trachomatis* (*Ctr*) is a bacterial pathogen with considerable medical importance. It is very common worldwide as an agent of sexually transmitted disease [1] and in some areas of the world relevant as the agent of ocular trachoma [2]. *Ctr* infects epithelial cells and, as an obligate intracellular bacterium, it depends on the integrity of the host cell. Apoptotic death of an infected cell has been noted many years ago as a defense mechanism against viral infection [3], and experimentally induced apoptosis disrupts the chlamydial developmental cycle as expected [4]. All tested species of the genus *Chlamydia* express an anti-apoptotic activity: cells infected with *Chlamydia* are profoundly protected against experimental pro-apoptotic stimuli, as first shown for infection with *Ctr* [5]. *Parachlamydia*, a *Chlamydia*-like organism that infects free-living amoebae, induces apoptosis and cannot grow in human or insect cells. When however the mitochondrial apoptosis pathway was disabled, growth of *Parachlamydia* in human cells was observed [6]; in insect cells, caspase inhibition had a similar effect [7]. Current knowledge therefore suggests that human cells can undergo apoptosis to defend the organism against the infection with *Chlamydia*, and human pathogenic chlamydial species had to evolve an anti-apoptotic strategy. Related *Chlamydia*-like species, whose hosts have no apoptosis system as in the case of *Parachlamydia* and amoeba, do not have the need of an anti-apoptotic mechanism and therefore cannot grow in human cells.

How the anti-apoptotic activity of *Chlamydia* works on a molecular level has been the focus of numerous studies, mostly using *Ctr*. *Ctr* induces many changes to cell signaling and metabolism, and a number of these changes have the potential to act in some anti-apoptotic way. However, the data point to a mechanism that operates at a central signaling step of the apoptotic pathway. The apoptosis signaling pathway is activated by numerous stimuli and converges on the activation of effector caspases, mostly caspase-3. Caspase-3 can be activated by caspase-9 in the mitochondrial pathway and by caspase-8 in the death receptor pathway [8]. Infection of human cells with *Ctr* blocks only mitochondrial apoptosis but not the death receptor pathway in the absence of a mitochondrial contribution [9]. In the mitochondrial pathway, the decisive step is the permeabilization of the outer mitochondrial membrane (MOMP), which leads to the release of intermembrane space proteins (in particular cytochrome *c*) that in the cytosol induce the activation of caspase-9 and the downstream steps of apoptosis. MOMP is regulated by the BCL-2 family of proteins. Within this family, one anti-apoptotic group (BCL-2 and its homologous ‘BCL-2-like’ proteins) directly bind and inhibit the two pro-apoptotic groups of proteins, BH3-only proteins and effector proteins (BAX and BAK). Upon their activation, BAX/BAK are the most downstream proteins of the pathway to MOMP, as they directly form pores in the outer mitochondrial membrane [10, 11].

Small molecule inhibitors of BCL-2-like proteins (so-called BH3-mimetics) have become available. At least in cell lines, they inhibit for instance BCL-2, which causes the direct activation of BAX/BAK, bypassing any upstream signals. This experimental approach has been used to map the anti-apoptotic activity of *Ctr*. As said above, there may be individual ways of apoptosis inhibition by *Ctr* infection in some pathways that also regulate apoptosis. The use of BH3-mimetics however identified a strong anti-apoptotic activity in *Ctr*-infected cells downstream of BCL-2-like proteins. This activity involved direct interaction of BAK with a protein generated during *Ctr* infection, which blocked BAK-activation (measured by conformational changes and oligomerization) at identifiable steps. The apoptotic activity of BAX, on the other hand, is in part counter-regulated through driving its retro-localization from mitochondria to the cytosol, and this activity was enhanced during infection, removing BAX from mitochondria and preventing its pro-apoptotic activity [12].

Intriguingly, molecularly similar anti-apoptotic activity has previously been described for a mitochondrial porin, VDAC2. Porins are β-barrel proteins inserted in the outer mitochondrial membrane or in the outer membrane of Gram-negative bacteria, where they regulate solute flux across the membrane. *Ctr* has a highly expressed porin, the major outer membrane porin (MOMP; we will here use the gene name OmpA to avoid confusion with mitochondrial outer membrane permeabilization). We expressed OmpA in (uninfected) HeLa cells and found that it localized to mitochondria where it inserted into the membrane. Furthermore, OmpA expression (in the absence of infection) precisely phenocopied the anti-apoptotic effect of *Ctr* infection on a molecular level [12]. This suggested that OmpA may be a major anti-apoptotic effector of chlamydial infection. This could be an attractive model: *Chlamydia*, as a Gram-negative bacterium, would use its evolutionary relationship to mitochondria, to inhibit apoptosis by mimicking the activity of the mitochondrial porin VDAC2.

The obvious difficulty with this model is the question how OmpA could translocate to mitochondria. *Ctr* replicates in the host cell within a cytoplasmic vacuole (the inclusion), so OmpA would have to be released from the bacteria, cross the inclusion membrane and traffic through the cell to reach mitochondria. However, immunostaining has identified OmpA outside the inclusion probably in vesicles, most likely outer membrane vesicles (OMVs) that appeared to move through the cell [13, 14]. Even though mechanistically unclear, OMVs from various bacterial species have been shown to traffic through mammalian cells and sometimes to target mitochondria, as shown for OMVs from *E. coli* [15] and from *Neisseria gonorrhoeae* [16]. We therefore here tested the hypothesis that OmpA indeed is a major anti-apoptotic factor during *Ctr* infection and reaches mitochondria through vesicular trafficking.

## Results

### Inhibition of apoptosis by bacterial porins

All tested *Chlamydia* species inhibit apoptosis. All of these have OmpA proteins that are closely related to *Ctr* OmpA. The only *Chlamydia*-like bacterium outside the *Chlamydiaceae* that is known to inhibit apoptosis is *Simkania negevensis* (*Sn*) (order *Simkaniales*) [17]. Intriguingly, OmpA genes in a non-*Chlamydiales* species in the data base are only found in *Sn* and in some not further characterized species of the *Simkaniales* and the *Rhabdochlamydia*. While lacking some regions of *Chlamydia* OmpA, *Sn*OmpA shows a high level of similarity to the OmpA from various *Chlamydia* species (Fig. S1A). We expressed *Sn*OmpA in human HeLa cells and tested for anti-apoptotic activity. As a control, we chose a random bacterial porin, OmpC from *E. coli* (*Ec*). As reported before [12], *Ctr*OmpA localized to the mitochondrial fraction of the cells to a substantial degree (Fig. 1A). *Sn*OmpA also showed strong mitochondrial localization while *Ec*OmpC mostly remained cytosolic (Fig. 1A). Expression of either *Ctr*OmpA or *Sn*OmpA protected the cells against apoptosis induced by BH3-mimetics, i.e. downstream of anti-apoptotic BCL-2-like proteins while *Ec*OmpC had no effect (Fig. 1B). AlphaFold predictions show that both *Ctr*OmpA and *Sn*OmpA have the typical β-barrel structure of porins (Fig. S1B). *Sn*OmpA is lacking certain stretches of *Ctr*OmpA, and these parts are in the loop regions outside the β-barrel structure (further discussed below). The identified anti-apoptotic activity of *Sn*OmpA, together with apoptosis inhibition by *Simkania negevensis* infection, is consistent with the hypothesis that OmpA is a relevant anti-apoptotic factor during infection with Chlamydia.

**Figure 1.**
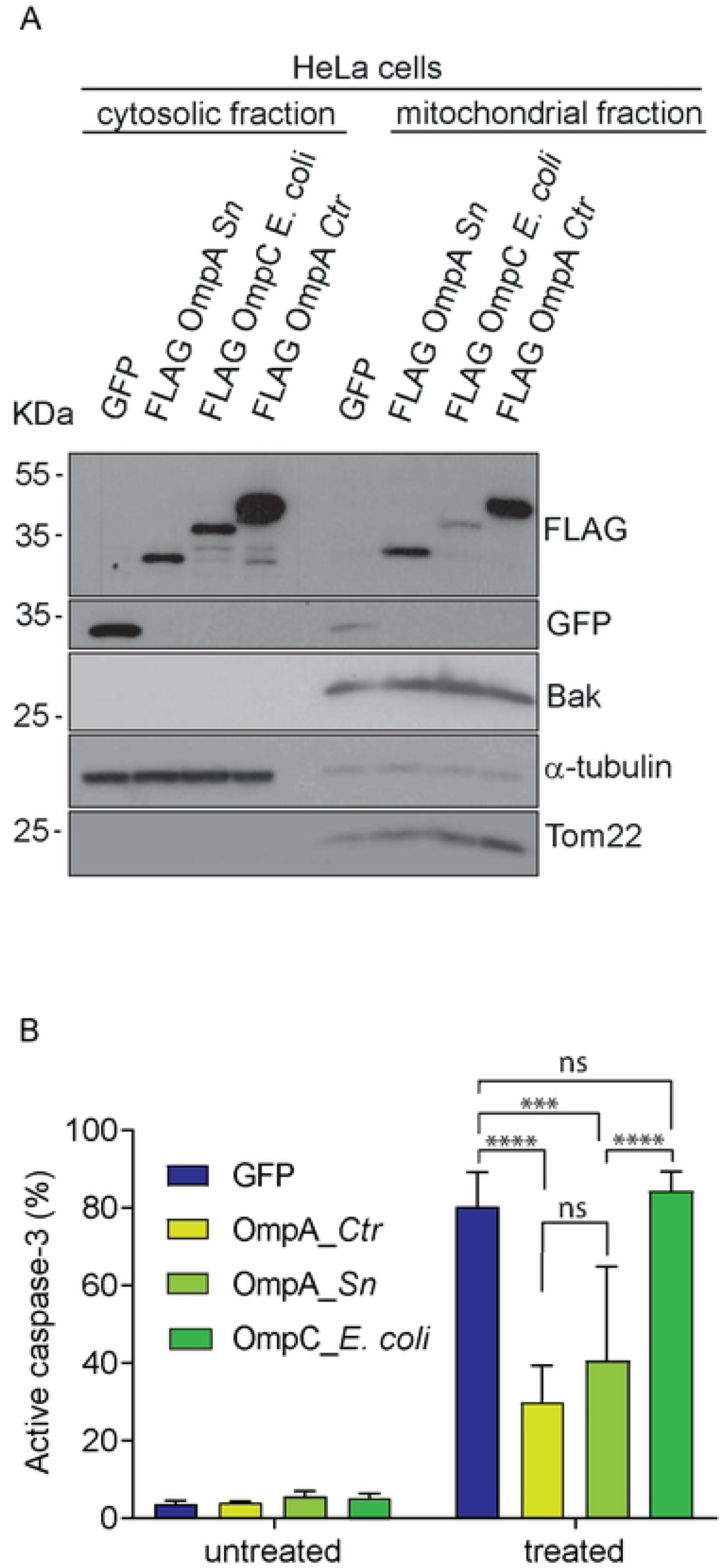
Mitochondrial localization of bacterial β-barrel proteins. **A**, HeLa cells constitutively expressing GFP, FLAG-OmpA from *Simkania negevensis* (*Sn*), FLAG-OmpC from *E. coli*, or FLAG-OmpA from *Ctr* were fractionated, and mitochondrial and cytosolic fractions were subjected to anti-FLAG or anti-GFP Western blotting to analyze the subcellular localization of GFP, OmpC or OmpA. Tom20 and Bak were used as mitochondrial markers, α-tubulin as a marker of the cytosol. **B**, *Sn*OmpA, *E. coli* OmpC and *Ctr* OmpA were ectopically expressed in HeLa cells. Cells were treated with ABT-737 (1 μM) and S63845 (500 nM) for 4h. Cells were fixed, permeabilized and stained for active caspase-3 to measure the number of apoptotic cells. Data are representative of three independent experiments. Error bars represent SEM and significance was tested using 2-way ANOVA (***, p<0.001, ****, p<0.0001).

### Interaction of ectopically expressed OmpA and BAK on mitochondria

We have previously reported that OmpA, when expressed in human cells in the absence of *Ctr* infection, inserts into mitochondrial membranes and can interact with BAK [12]. During activation, BAK undergoes a conformational change exposing its N-terminus [18]. At the time, the available anti-BAK antibodies only recognized active BAK, and we could show the interaction between OmpA and BAK only during the induction of apoptosis. More recently, an antibody against total BAK, including BAK in its inactive state, has become available [19]. Using this antibody, we found that OmpA bound to BAK also in its inactive state, in the absence of an apoptotic stimulus (Fig. 2A). This is similar to the constitutive interaction between BAK and VDAC2 and again supports the view that OmpA can replace VDAC2 in its BAK-inhibitory function. To test the BAK-OmpA interaction further we used proximity ligation assay (PLA). This assay utilizes antibodies to identify close interactions of the antibody targets inside cells (within a range of 30-40 nm) [20]. We detected a clear signal of the proximity of BAK and ectopically expressed OmpA (Fig. 2B, Fig. S2) on the mitochondria of cells, as shown by co-localization with the mitochondrial outer membrane protein TOM20 (Fig. 2C). These results confirm the finding that OmpA, when expressed in human cells, localizes to mitochondria and directly interacts with BAK in resting and activated state.

**Figure 2.**
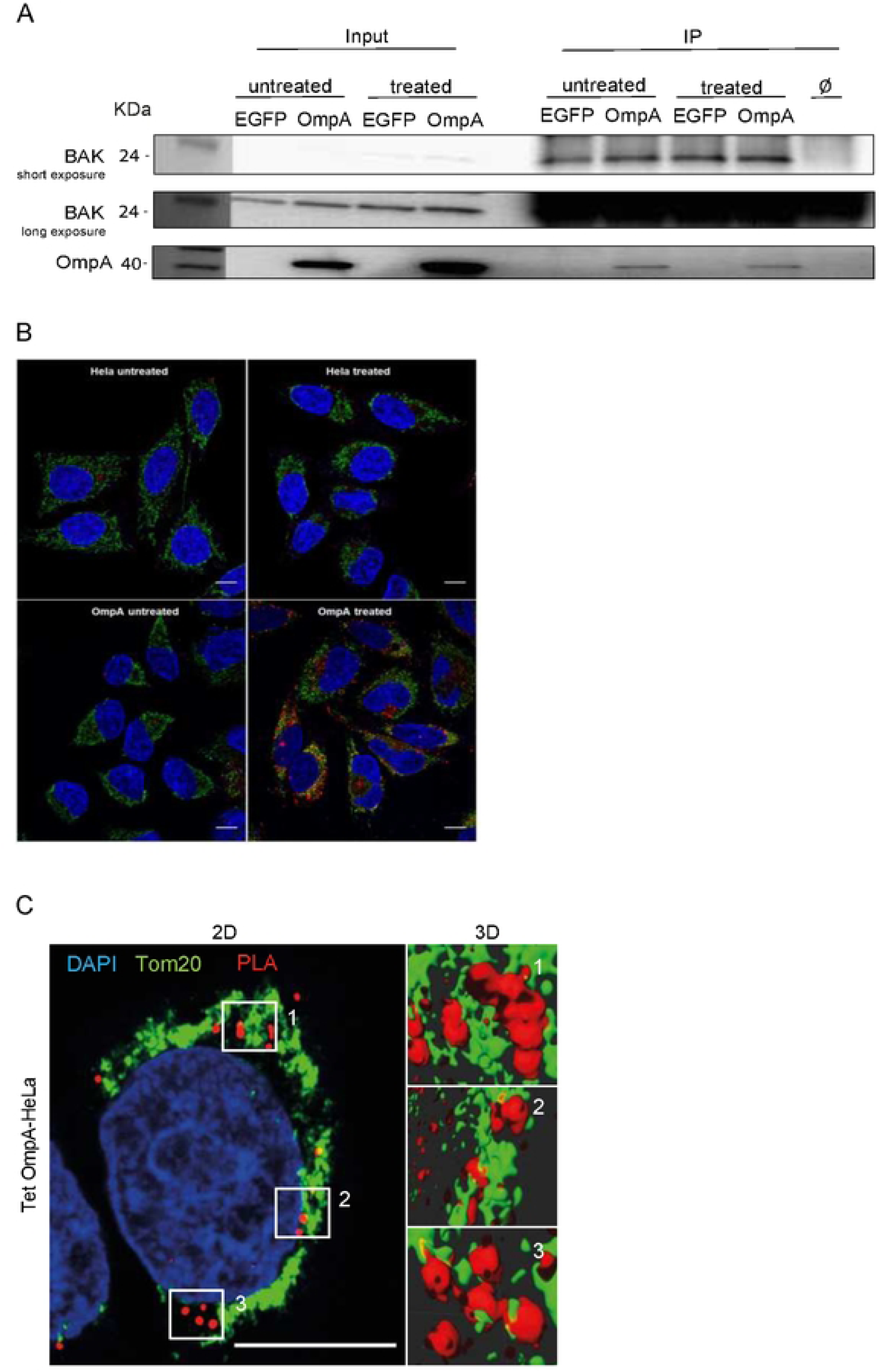
Interaction between OmpA and BAK on mitochondria. **A**, HeLa EGFP expressing cells and HeLa OmpA expressing cells were treated with ABT-737 (1µM) and S63845 (500nM) for 3h in the presence of the caspase-inhibitor QVD-OPh (10 μM; to block cell death downstream of mitochondria). BAK was immunoprecipitated with an antibody recognizing active and inactive BAK. Proteins were run on SDS-PAGE, and BAK and OmpA were detected by immunoblotting. Aliquots of the input and IP-reactions were loaded separately. Right lane shows beads with no antibody. Data are representative of three independent experiments. **B**, control HeLa cells (upper panel) and Tet-OmpA HeLa cells (carrying a tetracycline/AHT-inducible OmpA [12], bottom panel) were seeded on cover slips. 48h post-stimulation with AHT, cells were treated with ABT-737 (1 μM) and S63845 (500 nM) for 4h in the presence of the caspase-inhibitor QVD-OPh (10 μM). Cells were fixed, permeabilized and processed for PLA with antibodies against BAK (aa23-38; active BAK) and OmpA (red). Mitochondria were labeled using antibodies directed against Tom22 (green), and DNA was stained with Hoechst dye (blue). Confocal microscopy was performed and the overlay shows the co-localization of the PLA-signal with the mitochondrial protein Tom22. Images were acquired under identical conditions and exposure times. Data are representative of three independent experiments. Scaling bar, 10 μm. A control experiment with BAK-deficient cells is shown in Fig. S3. **C**, same set-up as in B. HeLa cells carrying a tetracycline-inducible OmpA were incubated with 100 nM AHT for 48 h to induce OmpA expression. Cells were fixed, permeabilized and processed for PLA using a different antibody specific for BAK (Ab-1(TC-100; active BAK)) and OmpA (red). Mitochondria were labeled using antibodies directed against Tom20 (green), and DNA was stained with Hoechst dye (blue). Cells were imaged by confocal microscopy. Stacks were then processed with the deconvolution software AutoQuantX and 3D reconstruction analysis was performed using the Imaris software. Images were acquired under identical conditions and exposure times. Left panel represents a single Z-slice, right panels indicate 3D reconstructions of the indicated areas using multiple Z-slices. Data are representative of three independent experiments. Scaling bar, 10 μm

### OmpA translocates to mitochondria during Ctr infection

To inhibit BAX/BAK, OmpA has to reach mitochondria during infection. We first tested whether the interaction of OmpA with BAK could be confirmed during infection in an unbiased proteomic screen of interaction partners. We infected HeLa cells with *Ctr* and treated them with an apoptotic stimulus (staurosporine) or not. For this screen, we used two different conformation-specific antibodies that both recognize only active BAK and should isolate BAK only when apoptosis is induced; this requirement for BAK activation therefore provides an additional specificity control for the immune-precipitation. We immuno-precipitated BAK and analyzed the isolated proteins by mass spectrometry (Fig. S3A). As expected, the amount of BAK detected was much greater upon apoptosis induction than in untreated cells (Fig. S3B). Although numerous proteins were found in the four analyses, the only protein that was consistently detected in all conditions was OmpA, and its identified levels were substantially greater in the samples from apoptotic cells (Fig. S3B). Western blotting of immuno-precipitates from *Ctr*-infected cells treated with BH3-mimetics confirmed the co-purification of OmpA with BAK during infection and apoptosis induction (Fig. S3C).

We then used single-molecule localization microscopy (SMLM) of infected cells to confirm the presence of OmpA outside the chlamydial inclusion and identified OmpA staining in TOM20 positive areas (i.e. mitochondria) (Fig. 3A, Fig. S4). Immuno-gold labelling and transmission electron microscopy further confirmed this finding: an OmpA signal was obtained in the cytosol and on mitochondria, consistent with the interpretation of a localization of OmpA on the outer mitochondrial membrane (Fig. 3B). We performed PLA of infected cells, again using antibodies against BAK and OmpA. As in OmpA-expressing cells, we found a clear PLA signal in close proximity of the mitochondrial protein TOM20 (Fig. 3C). Next, we isolated mitochondria from *Ctr*-infected cells. Following the fractionation of cells into cytosol and heavy membrane fraction (containing mitochondria), mitochondria were purified from this fraction using anti-TOM20 immuno-purification with antibody-labelled beads. Following this purification, we obtained mitochondria (identified by the mitochondrial membrane protein BAK and the mitochondrial intermembrane space protein SMAC), with no detectable ER (BIP) and with minimal contamination from nuclear membrane (lamin B) and endosomal compartments (Rab7). OmpA very clearly co-purified with mitochondria (Fig. 3D). To distinguish membrane insertion from other association with mitochondria, we extracted the mitochondria with sodium carbonate. This treatment removes any non-inserted, attached proteins before mitochondrial membranes are pelleted. OmpA was found exclusively in the membrane pellet while this disruption of mitochondria released the soluble intermembrane space protein SMAC into the supernatant as expected (Fig. 3D). These results show that during *Ctr*-infection OmpA translocates to mitochondria where it inserts into mitochondrial membranes.

**Figure 3.**
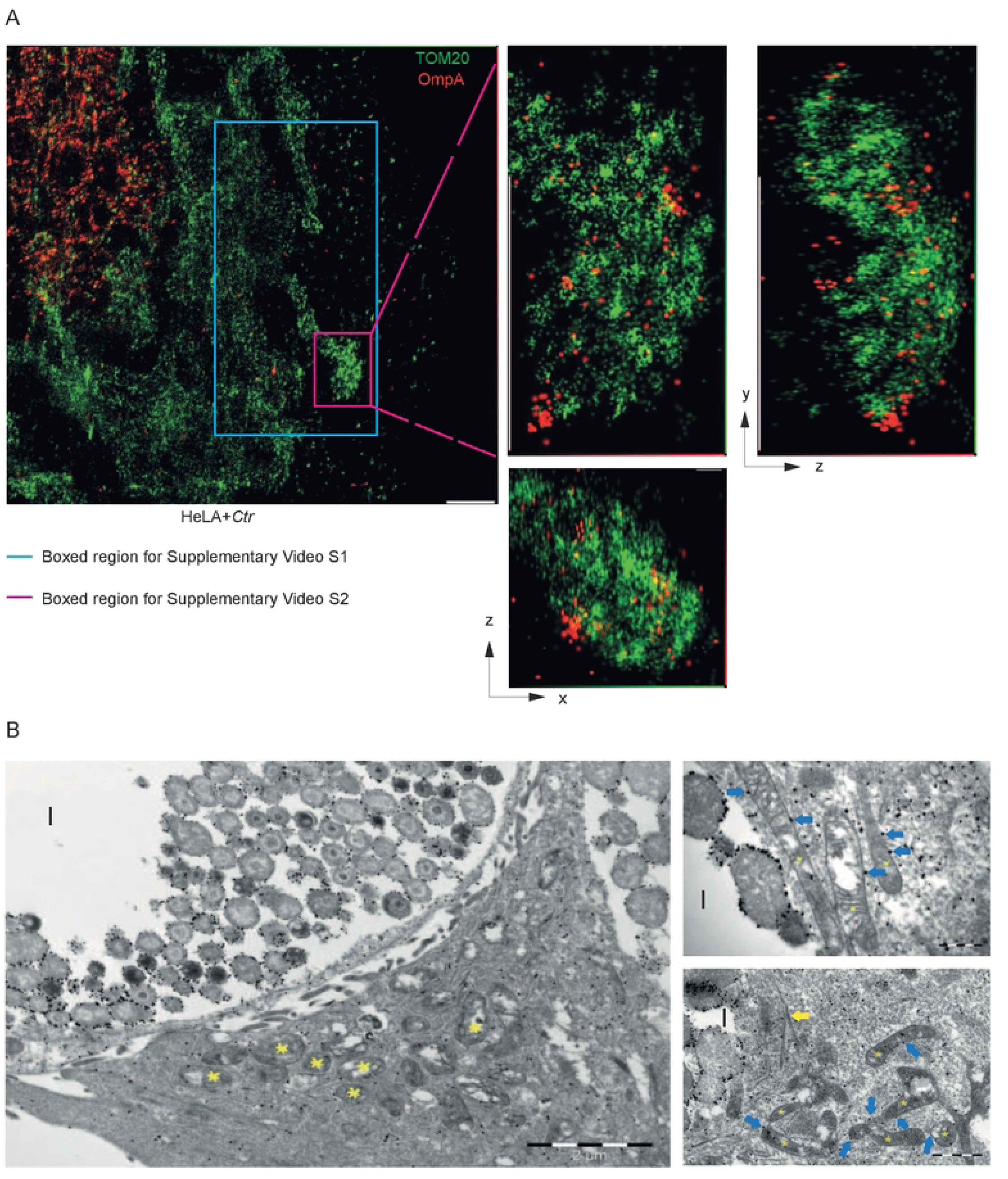
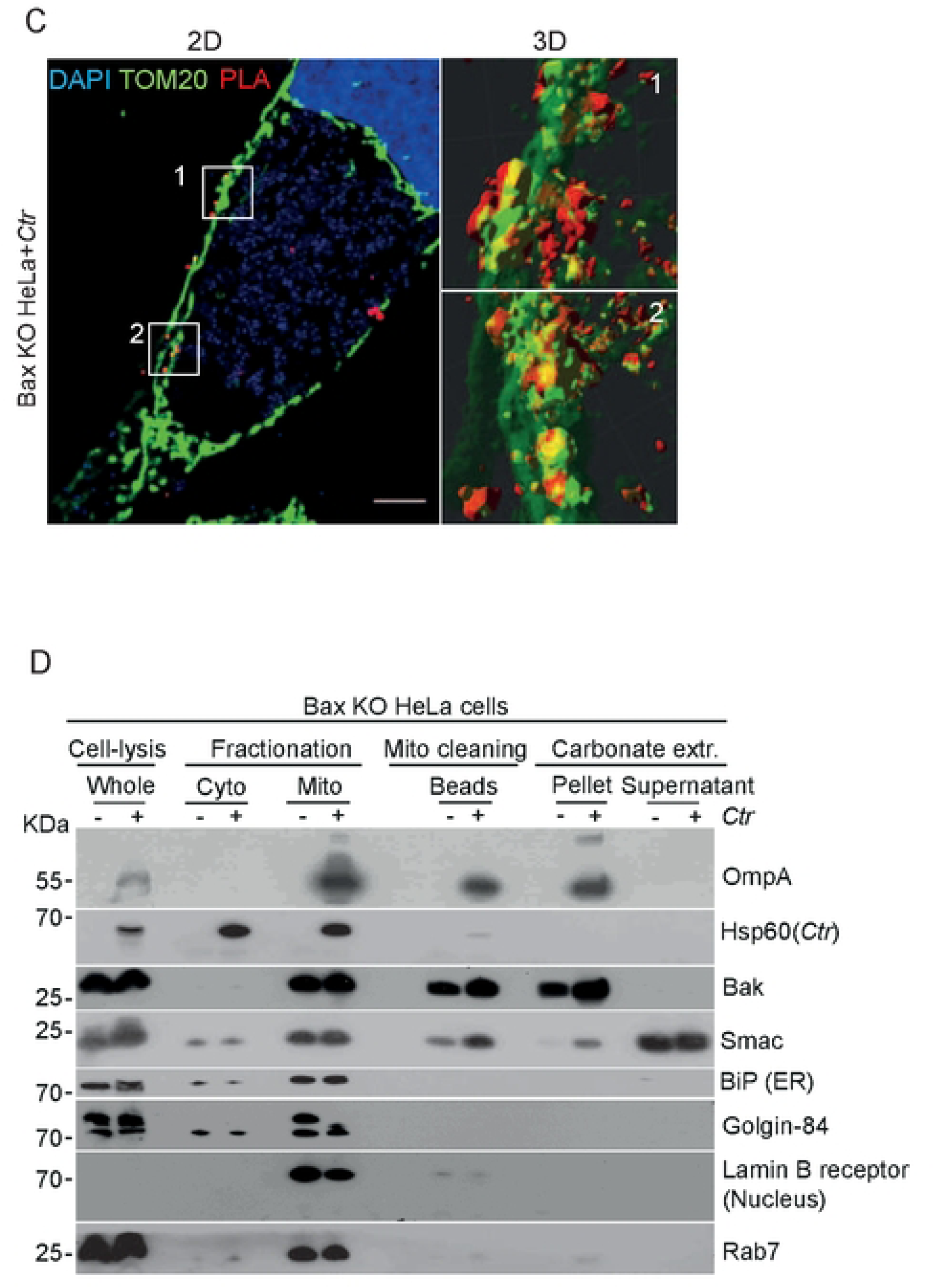
OmpA is detected at mitochondria during Ctr infection. **A**, 24 h after *Ctr*-infection, HeLa cells on cover slips were fixed, permeabilized and stained for two-color 3D SMLM super-resolution images of mitochondria and *Ctr* using antibodies against Tom20 and OmpA (left panel). The panels in the middle and to the right show magnified views of the pink boxed region. The x, y, and z-dimensions of the magnified region are 830 nm x 1490 nm x 780 nm. Scale bar: 1 μm. Note the localization of OmpA (red) on mitochondria Tom20 (green) during *Ctr*-infection. Insets show different perspectives of the pink boxed region. Also see Movies S1 and S2. **B**, immuno-gold electron microscopy identifies OmpA outside chlamydial inclusions and on mitochondria. HeLa cells were infected with *Ctr* and then fixed and stained for OmpA using immuno-gold particles and were imaged using electron microscopy as described in the methods. Yellow asterisks represent mitochondria, yellow arrows indicate cytoskeletal structures and blue arrows show gold particle labelling of OmpA on mitochondrial outer membranes. I (inclusion). Note the dense labeling of the bacteria as well as the labeling of mitochondrial membranes. Scale bar references are indicated. Data represents one experiment. **C**, BAX-deficient HeLa cells were seeded on cover slips. 24 h post-infection (MOI=5), cells were treated with ABT-737 (1 μM) and S63845 (500 nM) for 4 h in the presence of the caspase inhibitor QVD-OPh (10 μM). Cells were fixed, permeabilized and processed for PLA using antibodies against BAK (Ab-1(TC-100); active BAK) and OmpA. The red signal (PLA) indicates close proximity of BAK and OmpA. Mitochondria were stained using antibodies directed against Tom20 (green), and DNA was stained with Hoechst dye (blue). Cells were imaged by confocal microscopy. Stacks were then processed with the deconvolution software. AutoQuantX and 3D analysis was performed using the Imaris software. Left panel represents a single Z-slice, right panels indicate 3D reconstructions of the indicated areas using multiple Z-slices. Data are representative of three independent experiments. Scale bar, 5 μm. **D**, BAX-deficient HeLa cells were either *Ctr*-infected (MOI=5) or mock-infected. 24 h post-infection, cells were fractionated, and mitochondria were purified from heavy membrane fractions using magnetic beads labelled with anti-Tom20 antibodies. Purified mitochondria were subjected to sodium carbonate extraction (pH 11.5) to separate integral from attached membrane proteins. Membranes were pelleted, and the fractions were run on SDS-PAGE. Mitochondrial membranes and membrane-integrated proteins are found in the pellet fractions. Proteins were detected using the indicated antibodies. Chlamydial Hsp60, BiP (ER), Golgin-84 (Golgi-apparatus), lamin B (nuclear envelope) and Rab7 (endosomes) were used as organelle markers. Release of Smac shows extraction efficiency. Data are representative of three independent experiments.

### OmpA resides in membrane vesicles

The most likely way how OmpA could reach mitochondria seemed transport on vesicles, most likely OMVs. We used established OMV purification protocols to isolate vesicles from lysates of *Ctr*-infected cells. The purification yielded vesicular structures of about 150 nm (Fig. 4A). Although control isolations (the same procedure performed on lysates from uninfected cells) also yielded particular structures (Fig. S5), they were smaller and numbered about 10 % of the events from *Ctr*-infected cells. No vesicular structures could be detected by TEM on these control isolations (not shown). We will refer to these isolated structures as OMVs. OMVs from *Ctr*-infected cells contained OmpA but bacterial RNA polymerase B was not detectable, indicating that no bacterial particles had been co-purified. Likewise, no mitochondrial VDAC was detectable (Fig. 4B).

**Figure 4.**
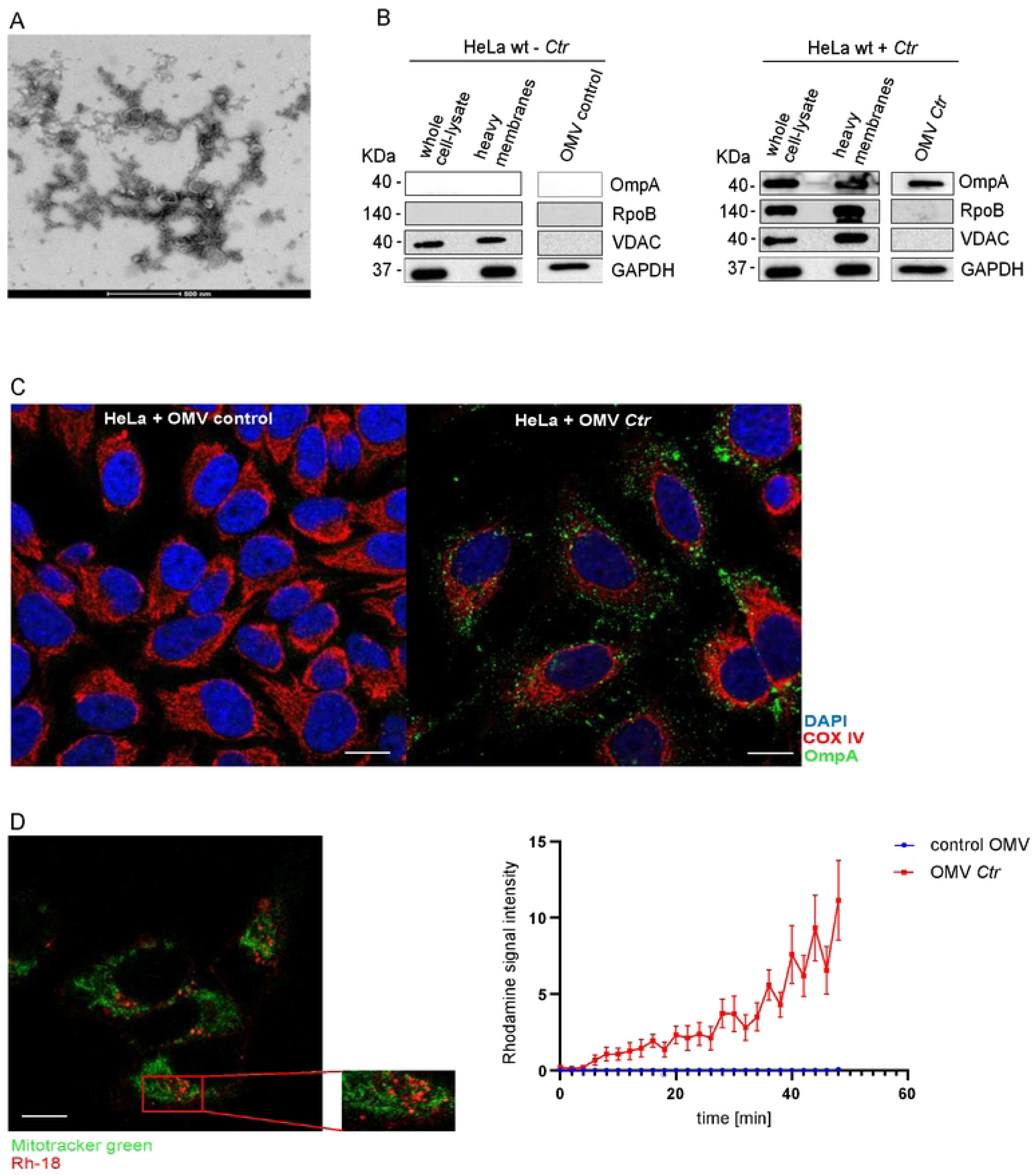
OmpA resides in membrane vesicles. **A**, electron microscopy was performed on chlamydial outer membrane vesicles, which were isolated from *Ctr*-infected HeLa cells. **B**, OMVs were isolated from *Ctr*-infected (MOI=5, 48h) HeLa cells. Uninfected cells were subjected to the same procedure. Isolated OMV-containing fractions were subjected to Western blotting. OmpA but not the chlamydial RNA polymerase B (RpoB, as a marker for bacteria) was seen in vesicles purified from *Ctr*-infected HeLa cells. VDAC was used to detect mitochondrial contamination. GAPDH was used as a loading control. Data are representative of three independent experiments. **C**, HeLa cells were seeded on cover slips. Cells were incubated with chlamydial OMVs or fractions isolated from uninfected control cells. Cells were treated with ABT-737 (1 μM) and S63845 (500 nM) in the presence of the caspase-inhibitor QVD-OPh for 3h. Cells were fixed and permeabilized. Mitochondria were labeled using antibodies directed against COX IV (red, inner mitochondrial membrane), OmpA (green), and DNA was stained with DAPI (blue). Confocal microscopy was performed with Zeiss LSM880 and the overlay shows the co-localization of the OMVs with the mitochondrial protein COX IV. Images were acquired under identical conditions and exposure times. Data are representative of three independent experiments. Scale bar, 10 μm. **D**, OMVs were labelled with rhodamine B chloride (R-18) (red) and were added to uninfected HeLa cells. Live cell imaging was performed for 50min on the confocal microscope Zeiss LSM 880. Mitotracker green was used to stain mitochondria. Graph displays fusion events of the R-18-labelled OMVs with host cell membrane over time. Scale bar, 10 μm. Data are means/SEM from three independent experiments. Control OMVs refers to fractions isolated from uninfected Hela cells.

The ability of OMVs from other bacteria to enter mammalian cells has been mentioned above. When we added *Ctr* OMVs to uninfected HeLa cells, an OmpA signal was detectable by confocal microscopy near mitochondria (Fig. 4C), suggesting that OmpA was transported on the OMVs into the cell. We tested whether the OMVs may be able to fuse with host membranes. For this purpose, we labelled isolated OMVs with rhodamine-R18 dye. At high concentrations, rhodamine-R18 is quenched in the vesicular membrane but dilution of the dye – which occurs upon fusion with other unlabeled membranes – de-quenches it, causing red fluorescence [21]. We added labelled OMVs to uninfected HeLa cells and observed a substantial number of fusion events over 50 minutes of recording by microscopy (Fig. 4D). Staining of TOM20 was consistent with the view that OMVs fused with mitochondria (Fig. 4D, bottom).

### OMVs deliver OmpA to BAK and inhibit apoptosis

We used PLA again to test for proximity of mitochondrial BAK and OmpA delivered by OMV when they were added to uninfected cells. We obtained a signal that was surprising in its clarity: in the conditions used, about 50 PLA signals could be detected per individual cell (Fig. 5A). When we isolated heavy membrane fractions from these cells, OmpA seemed to be restricted to the membrane/mitochondrial fraction, supporting the concept of mitochondrial fusion of the OMVs (Fig. 5B). The apoptosis effector protein BAX constantly diffuses from the cytosol to mitochondria and is retro-translocated to the cytosol by BCL-X_L_ [22] and by VDAC2 [23] as an anti-apoptotic regulatory mechanism. As reported previously, expression of OmpA in human cells also causes enhanced BAX retro-translocation, similar to the effect observed during *Ctr* infection [12]. As shown in Fig. 5B, addition of OMVs from *Ctr*-infected cells also caused a redistribution of BAX from mitochondria to the cytosol, suggesting an effect of OmpA on mitochondrial BAX. We finally tested whether the addition of OMVs to uninfected cells could inhibit apoptosis. A significant reduction of BH3-mimetic-induced apoptosis by OMV addition was indeed seen (Fig. 5C). Because of the effect on BAX retro-translocation, we also tested for apoptosis inhibition in BAK-deficient cells (where apoptosis relies exclusively on BAX). In these cells, the anti-apoptotic effect of OMV-addition was very strong (Fig. 5C). The results suggest that OmpA can travel on OMVs, insert into mitochondria through membrane fusion and inhibit apoptosis.

**Figure 5.**
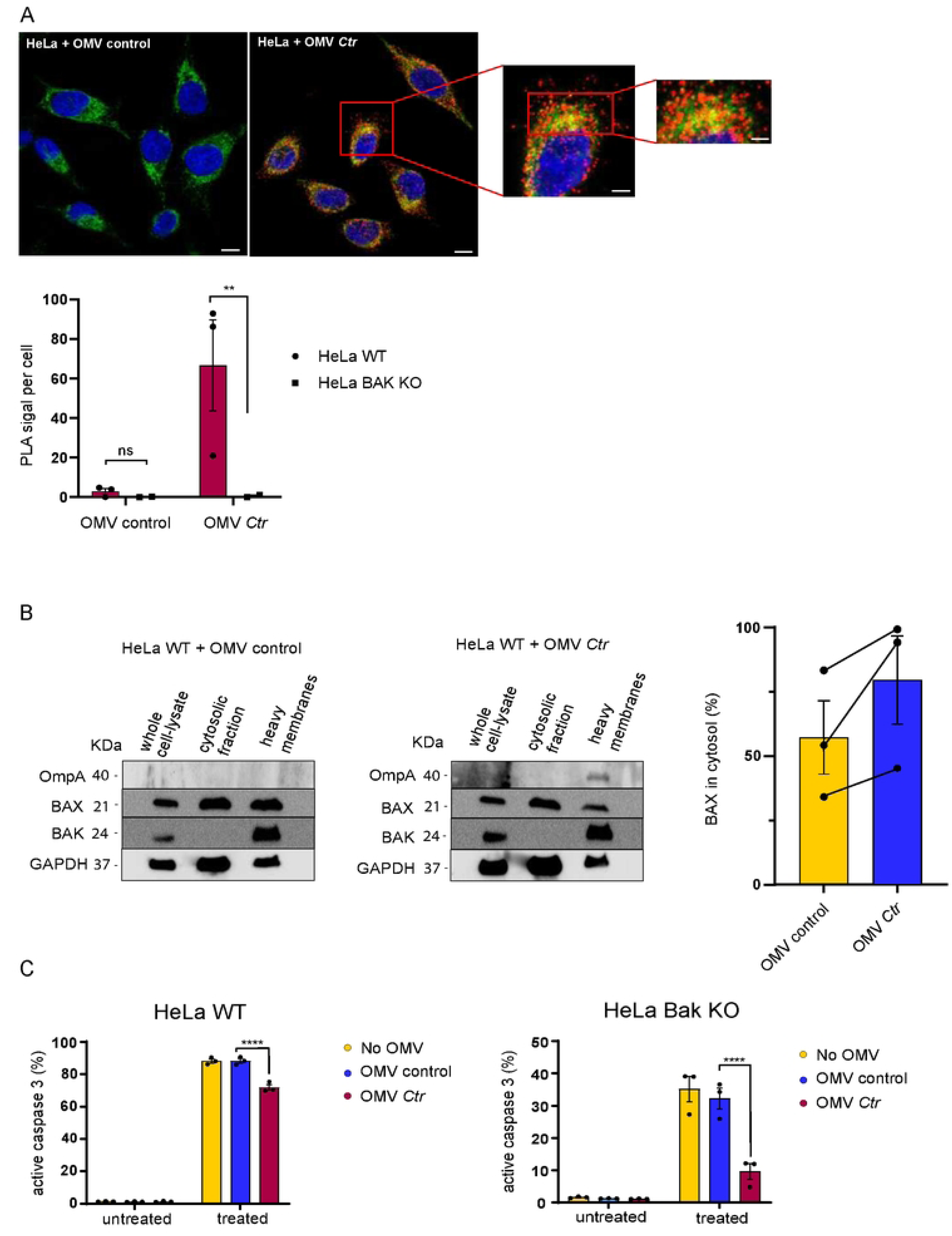
OMVs deliver OmpA to BAK and inhibit apoptosis. **A**, HeLa cells were seeded on cover slips. Cells were incubated with either chlamydial OMVs or preparations from uninfected HeLa cells (OMV control) for 1h. Cells were then treated with ABT-737 (1 μM) and S63845 (500 nM) for 3h in the presence of the caspase-inhibitor QVD-OPh (10 μM). Cells were fixed, permeabilized and processed for PLA using antibodies against active BAK (Ab-1(TC-100)) and OmpA (red) (Biozol Diagnostica, #LS-C79219). Mitochondria were labeled using antibodies directed against Tom20 (green) and DNA was stained with DAPI (blue). Confocal microscopy was performed and the overlay shows the co-localization of the PLA signal with the mitochondrial protein Tom20. Images were acquired under identical conditions and exposure times with the confocal microscope Zeiss LSM 880. BAK-deficient cells were used as a specificity control. Scale bar, 10 μm. The diagram shows the quantification from three independent experiments (columns are mean/SEM). Significance was calculated using the Kolmogorov–Smirnov test (**, p<0.01). **B**, HeLa cells were treated with chlamydial OMVs of fractions isolated from uninfected cells (control OMVs) for 2h. Cytosolic and mitochondria-containing (heavy membrane) fractions were separated and Western blot analysis was performed. OmpA is found on the heavy membrane fraction on HeLa cells treated with chlamydial OMVs. BAX retro-translocation to the cytosol is enhanced upon infection. Columns/error bars give means/SEM of three individual experiments (individual results are shown as symbols). GAPDH was used for normalization. Lines connect results of the experiments. Data are representative of three independent experiments. **C**. HeLa cells (control or BAK-deficient cells) were treated with chlamydial OMVs or fractions isolated from uninfected cells (control OMVs) for 1h. Cells were then treated with ABT-737 (1 μM) and S63845 (500 nM) for 3h. Cells were fixed, permeabilized and active caspase-3 staining was performed to measure the number of apoptotic cells. Data are means/SEM of three individual experiments. Significance was tested using 2-way ANOVA (****, p<0.0001).

## Discussion

We had previously reported that OmpA, when expressed without infection, could inhibit BAK and increase the retro-translocation of BAX to the cytosol. Because these molecular effects were exactly the same as the anti-apoptotic effects identified during *Ctr* infection [12], the results suggested that OmpA may be a mediator of the anti-apoptotic effect of chlamydial infection. How OmpA may translocate to mitochondria was unclear. Here we provide evidence that OmpA during infection reaches mitochondria on OMVs. This supports the concept that *Chlamydia* indeed uses its relationship with mitochondria and has evolved its porin OmpA to function as an apoptosis inhibitor, assuming the molecular function of the mitochondrial porin VDAC2.

The porins from various bacteria can insert into mitochondria *in vitro*, as shown for Neisseria PorB [24, 25], *Yersinia* YadA [26] or *E. coli* PhoE [27]. This indicates a degree of conservation between mitochondrial proteins and their bacterial ancestors. The porin from *Neisseria gonorrhoeae*, PorB, has been identified on mitochondria of infected cells, where it induced apoptosis [24], and this translocation has been suggested to occur on OMVs [16]. Surprisingly, purified PorB from *Neisseria meningitidis*, conversely, has been described to inhibit apoptosis. In that study, recombinant protein was added to human cells [28] but it is unclear how the protein would cross membranes. The ability of bacterial OMVs to enter human cells from the extracellular space is however well documented, and a number of uptake mechanisms have been proposed [29]. Mitochondrial targeting of OMVs has further been described: *E. coli* OMVs can deliver the hemolysin HlyA to mitochondria where it induced apoptosis [15]. When OMVs from *N. gonorrhoeae* were incubated with macrophages, they induced apoptosis and PorB co-localized with TOM20. Because experimental PorB expression induced apoptosis, the authors concluded that PorB on OMVs induced apoptosis in that situation. OMVs from *Acinetobacter baumannii* induced mitochondrial fragmentation, although this depended on the GTPase DRP1 and may therefore not reflect direct mitochondrial targeting of OMVs [30]. Why OMVs target mitochondria is not obvious. It is clear that chlamydial OMVs fuse with mitochondrial membranes but there may be fusion with other intracellular membranes as well. One possibility of how specificity could be achieved is the physical interaction between OmpA and BAK, which is inserted into the outer mitochondrial membrane; BAK recognition by OmpA may represent a targeting mechanism. It may also be an effect of different lipid composition of the intracellular, organellar membranes.

Our results show fusion events of OMVs with the mitochondrial membrane. OmpA is inserted into the mitochondrial membrane during *Ctr* infection, as shown by carbonate extraction (Fig. 3D). When the membrane of OMVs fuses with the mitochondrial outer membrane, this can be expected directly to transfer OMV-membrane inserted OmpA into the mitochondrial membrane. A number of aspects are still unclear – how do the OMVs cross the inclusion membrane? When they fuse with the mitochondrial membrane, what will be the orientation of OmpA? To approach the question of BAK interaction, we modelled the complex of β-barrel proteins and BAK using AlphaFold. Experimental mutagenesis and modelling has yielded a structural model of the interaction of BAK and VDAC2, with BAK sitting ‘on top’ of membrane-inserted VDAC2, and the BAK transmembrane domain stretching alongside VDAC2. The AlphaFold model of the complex (Fig. S6A) is very similar to this in part experimentally confirmed model [31]. Intriguingly, the AlphaFold model of OmpA and BAK shows core similarities. BAK is placed similarly on top of the β-barrel, with plausible direct interaction between the proteins. Further, the BAK C-terminal transmembrane helix also extends into the membrane alongside the β-sheets (Fig. S6B; AlphaFold2 cannot model membrane interactions so this structure was predicted purely for the proteins). The sequence alignment of the various OmpA proteins identified a strand-loop-strand region that was conserved between *Ctr*OmpA and *Sn*OmpA (Fig. S1A, last two beta strands of alignment). In the predicted structure, this loop aligns with the transmembrane helix of BAK and may provide direct binding (Fig. S6A). To test this hypothesis, we mutated the motif in *Ctr*OmpA but were unfortunately unable to achieve expression of the mutated construct in HeLa cells, even following codon optimization (not shown). OmpA has been proposed to exist as a trimer. Modelling a trimeric OmpA complex with three BAK molecules in AlphaFold shows a different orientation of the barrels in relation to BAK, but the alignment of the BAK transmembrane helix to the barrel of OmpA remains (Fig. S6C). We cannot of course confirm which of these models is more accurate at the moment.

The orientation of the binding in the model is noteworthy: based on this prediction, the extracellular regions of OmpA protrude into the mitochondrial intermembrane space, and the periplasmic regions will become cytosolic. This suggests that the fusion of OMVs is such that the orientation of OmpA is reversed compared to its orientation in the outer bacterial membrane. A number of ways have been proposed how OMVs fuse with human membranes, through lipid mixing or through protein interaction [29]. It is possible that the vesicles already flip over and invert when moving across the inclusion membrane; the orientation of OmpA as the vesicles fuse with mitochondria is uncertain. The predicted model of OmpA-BAK protein interaction seems so plausible and so similar to VDAC2-BAK that we think it has a high likelihood of being correct.

Most bacteria are not obligate intracellular parasites and apoptosis has a minor function in the interaction of human cells and bacteria [32]. *Chlamydia* is an exception, in that it can only grow inside host cells and very likely has to inhibit apoptosis to be able to grow [6, 33]. The principle of apoptosis inhibition is much clearer for viruses where anti-apoptotic activities have been known for a long time [34]. It is also easier for viruses to come up with anti-apoptotic strategies: large DNA viruses, for instance, often carry genes whose products are similar to anti-apoptotic BCL-2-like proteins [34]; the viruses have acquired host cell genes and modified them for their purpose. Acquisition of genes from mammalian cells is not as straightforward for bacteria, and there is no evidence of horizontal transfer of mammalian genes to *Chlamydia*. We find it very interesting that *Chlamydia* appears to exploit its evolutionary relationship with mitochondria in this way. Mitochondria have acquired a number of functions in their host cell. While other apoptosis regulators are probably more important – mainly the BCL-2 family – there is a clear function in the regulation of apoptosis for VDAC2 and its interaction with BAX and BAK. *Chlamydia* appears to utilize its OmpA to inhibit apoptosis in a way that is probably very similar to the interaction of BAK with VDAC2. While OmpA has very likely essential functions in regulating solute flux across the membrane, its second function in inhibiting apoptosis may also be a great relevance.

## Methods

### Cell lines and cell culture conditions

HeLa 229 cervical carcinoma cells (ATCC) and 293FT cells (Invitrogen) were maintained in RPMI medium (Life Technologies, UK) supplemented with 10% FCS (tetracycline negative; Life Technologies).

### Construct and generation of cell lines

OmpC was cloned from E. coli (NEB, #C3040H). *Simkania negevensis* (*Simk*) genomic DNA was kindly provided by Dr Matthias Horn, Vienna. OmpA from *Simk* and OmpC from *E. coli* ORFs without their respective signal peptides were amplified by PCR using the following primers: for *Simkania* (Flag-OmpA (*Simk*)_del1-18-GW-Sense; 5’-caccATGGATTACAAGGATGACGATGACAAGTTGTATAACGGCAATCCAAGT-3’ and OmpA (*Simk*)-antisense; 5’-CTAGAACTTCACTTCGCCG-3’) and for E. coli (Flag-OmpC_del1-21-GW-Sense;5’-caccATGGATTACAAGGATGACGATGACAAGGCTGAAGTTTACAACAAAGAC-3’ and OmpC-antisense; 5’-TTAGAACTGGTAAACCAGACC-3’). The ORFs (Gene ID: 946716 for OmpC (*E. coli*) and OmpA (*Simk*), SNE_A00410 were subsequently cloned into pENTR/SD/D-TOPO Gateway vector (Life Technologies) and were then inserted into the lentiviral vector pFCMVTO_GW_SV40_PURO_W via Gateway LR recombinase reaction (Life Technologies). Lentiviruses were used to infect HeLa cells to establish cell lines stably expressing FLAG-OmpA (*Simk*) or FLAG-OmpC. For inducible OmpA, HeLa cells were first transduced with lentivirus expressing Tet repressor, and selected cells were then infected with lentivirus carrying untagged OmpA. HeLa cells expressing either inducible or constitutive GFP were generated as above by cloning GFP into pFCMVTO_GW_SV40_PURO_W. Cells stably carrying the construct were selected with 1 μg/ml of puromycin (InvivoGen, #ant-pr-1).

### Lentivirus production and bacterial infection

For lentivirus production, 293FT cells were transfected with corresponding vectors using FuGene HD transfection (Promega, USA), following the manufacturer’s instructions. Packaging vectors were psPAX.2 and psMD2.G (Addgene Plasmids, #12260 and #12259; Didier Trono). Virus-containing supernatant was collected, filtered and incubated with HeLa cells in the presence of 1 μg/ml of polybrene (Millipore, #TR-1003-G). *Chlamydia trachomatis* serovar L2 (*Ctr*) was obtained from ATCC and propagated in HeLa cells. Bacteria were purified over a Gastrografin density gradient (Bayer Vital, Leverkusen), followed by titration on HeLa cells and stored in SPG medium (0.2 M sucrose, 8.6 mM Na2HPO4, 3.8 mM KH2PO4, 5 mM glutamic acid [pH 7.4]) at 80°C. Fresh aliquots were thawed for each experiment. Cells were infected at a multiplicity of infection of 5 in complete culture medium without antibiotics.

### Analysis of apoptosis by flow cytometry

Apoptosis was induced by treatment with the combination of ABT-737 (Selleck Chemicals, #S1002) and the Mcl-1-inhibitor S63845 (APExBio # A8737). Cells were collected, washed and fixed with 4% paraformaldehyde (Morphisto, #11762.00250) for 10 min at room temperature, followed by staining with anti-active caspase-3 antibody (BD Pharmingen, #559565) in PBS (Life Technologies, #14190169) containing 0.5% Saponin (Roth, #4185.1) and 0.5% bovine serum albumin (BIOMOL, #BSA-50) for 30min at room temperature. Alexa Fluor 647-conjugated donkey anti-rabbit IgG (Dianova, #711605152) was used as secondary antibody and cells were analyzed by flow cytometry using a FACS Calibur Flow Cytometer (Becton-Dickinson, Heidelberg).

### Mitochondria isolation and determination of membrane insertion using sodium carbonate-extraction

Cells were harvested, washed and resuspended in MB-EDTA buffer. Mitochondria were isolated by passing cells through a 27G needle using 1 mL syringe. 24 h post-infection, cells were fractionated and mitochondria were purified from heavy membrane fractions using magnetic beads conjugated to TOM20 (SantaCruz, #sc-11415). Cleaned mitochondria were subjected to sodium carbonate extraction (pH 11.5) to separate integral from attached membrane proteins. Fractions were boiled in Laemmli-buffer at 95°C and run on SDS-PAGE. Mitochondrial membranes and membrane-intergrated proteins) are found in the pellet fractions. Proteins were detected using the indicated antibodies. Chlamydial OmpA and Hsp60, BiP (ER), Bak, Golgin-84 (Golgi-apparatus), lamin B receptor (nuclear envelope) and Rab7 (endosomes) were used as organelle markers. Release of Smac shows extraction efficiency.

### Immunochemistry and confocal microscopy

For OmpA staining, cells were seeded on coverslips and treated as indicated. After fixing for 15 min in 4% PFA, cells were permeabilized for 10 min in 0.2% Triton X-100 (Sigma) in PBS, and incubated for an additional 30 min in 5% BSA in PBS. Polyclonal goat anti-OmpA (Biozol Diagnostica, #LS-C79219) was used against OmpA, followed with Alexa Fluor 488-conjugated donkey anti-goat IgG (Thermo Fisher Scientific, #A-11055). To investigate mitochondrial localization of OmpA, mitochondrial protein TOM20 was co-stained with OmpA using monoclonal Mouse anti-TOM20 (SantaCruz, #sc-11415), followed by Cy5-conjugated donkey anti-mouse IgG (Dianova, #715-175-151). DNA was stained with DAPI (2 mg/ml; Sigma, #D9542) for 10 min before being mounted in Permafluor (Thermo Fisher). Images were taken using a Zeiss LSM 880 confocal microscope at a 63x magnification (oil immersion) and analyzed with ZEN 3.0 (Zeiss) software.

### Proximity Ligation Assay

Cells were seeded on coverslip and infected with *Ctr* or incubated for 1h with control OMVs or chlamydial OMVs. Cells were then treated with 1µM ABT-737 (Selleck Chemicals, #S1002) and 500nM Mcl-1-inhibitor S63845 (APExBio # A8737) for 2.5h in the presence of caspase inhibitor QVD-OPh (Gentaur (ApexBio), #GEN2269261) 10µM. Cells were washed and fixed with 4% PFA for 10min at room temperature. Mitochondria were first stained with polyclonal rabbit anti-Tom20 antibody (SantaCruz, #sc-11415), followed by Alexa Fluor 488-conjugated donkey anti-rabbit IgG (Dianova, #711-545-152). Duolink Proximity Ligation Assay (Sigma, #DUO92006-30RXN) was performed according to the manufactureŕs instructions. Antibodies used were monoclonal mouse anti-Bak (Millipore, #AM03) and polyclonal goat anti-OmpA (Biozol Diagnostica, #LS-C79219), followed by labelling with the corresponding secondary antibodies. After DNA staining with DAPI (Sigma, #D9542), images were taken using a Zeiss LSM 880 confocal microscope at a 63x magnification (oil immersion) and analyzed with ZEN 3.0 (Zeiss) software.

### Electron Microscopy of outer membrane vesicles

Cell lysates were fixed with 4% paraformaldehyde in 0.025 M cacodylate buffer (pH 7.4) for 1 hour at room temperature. For negative staining, 3.5 µL of the fixed sample was applied to glow-discharged 300-mesh carbon-coated copper grids (Science Service, catalog #ECF300-Cu-50) and allowed to adsorb for 60 seconds. Excess liquid was gently blotted with filter paper, and the grids were washed twice with ultrapure water. The samples were then negatively stained by applying a 10 µL drop of 2% uranyl acetate for 30 seconds, followed by blotting to remove excess stain. After air-drying at room temperature, the grids were examined using a Talos L120C transmission electron microscope (ThermoFisher Scientific) operated at 120 kV.

### Immuno-Gold Electron Microscopy

Cells were infected with *Ctr* (MOI=5) for 48h before they were fixed for 30 min at room temperature in 4% paraformaldehyde plus 0.05% glutaraldehyde in 0.1 M phosphate buffer. Afterwards, cells were permeabilized with 0.1% Triton X-100 (Sigma) for 3min, quenched with 0.05% glycine for 5min and blocked in 5% bovine serum albumin (BIOMOL, #BSA-50) for 30min. For immune-gold labelling cells were incubated with primary goat anti-OmpA antibody (Biozol Diagnostica, #LS-C79219) in 1% BSA at a dilution of 1:500 for 2h at room temperature. Washing steps were followed by incubation in secondary antibody coupled to 1.4 nm gold (Nanogold-Fab’ rabbit anti-goat IgG, Nanoprobes, USA, #2006) in 1% BSA at a dilution of 1:100 for 2h at room temperature. Several washes were performed before the post fixation buffer with 4% PFA + 2% Glutaraldehyde in 0.1M Sodium Cacodylate Buffer pH 7.4 was added to the cells for 15min at room temperature. Lastly, nanogold was enhanced for exactly 8 minutes using silver enhancement Kit (HQ silver, Nanoprobes, USA) in complete darkness. Cells were contrasted in 1% osmium tetroxid and 1 % uranyl acetate (in 70% ethanol) both for 30 min at RT. After dehydration, cells were embedded in epoxy resin (Durcupan, Sigma Aldrich) and ultrathin sections were cut using a Leica UC6 ultramicrotome. For imaging a Zeiss TEM 910 was used.

### Rhodamine-18 membrane fusion

OMVs were pelleted at 120,000xg for 30min at 4°C and resuspended in labeling buffer (50 mM Na2CO3, 100 mM NaCl, pH 9.2). Rhodamine isothiocyanate B-R18 (Molecular Probes) was added to the OMVs at a concentration of 1 mg/ml for 1 hour at 25°C, 300rpm followed by ultracentrifugation at 120,000 g for 30 min at 4°C. Rhodamine labeled-OMV were resuspended in PBS (0.2 M NaCl) and pelleted again. Labeled-OMVs were resuspended in 1 ml PBS (0.2 M NaCl) containing a protease inhibitor cocktail tablet (Complete Protease Inhibitor Tablet, Roche). R-18 labeled OMVs were added to HeLa cells. Live cell imaging was performed for 50min on the confocal microscope Zeiss LSM 880. Mitotracker green was used to stain mitochondria.

### BAK immunoprecipitation

Bak protein was immunoprecipitated as described [35]. Cells were treated with 1µM ABT-737 and 500nM Mcl-1-inhibitor S63845 for 3h in the presence of caspase inhibitor QVD-OPh (Gentaur (ApexBio), #GEN2269261) (10 µM). Cells were harvested and directly lysed in buffer containing 1% CHAPS (Carl Roth, #1479.2). Following centrifugation, protein concentration of the supernatants was quantified using Bradford assay (Bio-Rad) and equal amounts of protein lysates were loaded on the gel. Anti-BAK antibody (either clone 7D10; Walter and Eliza Hall Institute, Melbourne, Monoclonal Antibody Facility, total BAK, or clone aa23-38, active BAK) was incubated with protein G agarose beads (Roche, #11719416001) for 30min at 4°C to allow the antibody to bind to the beads. Afterwards, lysates were incubated with anti-BAK antibody and protein G agarose beads overnight at 4°C under constant agitation. Samples were collected, boiled in Laemmli buffer containing DTT for 5min at 95°C, separated by SDS-PAGE and immunoblotted.

### Nanoparticle tracking analysis

The concentration and size distribution of OMV were measured using the Nanoparticle Tracking Analysis (NTA) instrument Zetaview QUATT (Particle Metrix, Meerbusch, Germany) using the software ZetaView 8.4.2. For the experiments, the instrument was calibrated as described earlier [36]. Each sample was diluted tenfold in PBS, with the number of events between 70 and 200 events/frame. For determining the size distribution, each sample was measured at 11 different positions (5 cycles) in technical duplicates.

### Preparation of samples for proteomic analysis

For stable isotopic labeling by amino acids in cell culture (SILAC), cells were labeled with either L-arginine (Arg0) and L-lysine (Lys0) (‘light’ amino acids) or with 13C615N4 L-arginine (Arg10) and 13C615N2 L-lysine (Lys8) (‘heavy’ amino acids) (Silantes, #282986444) in DMEM supplemented with 10 % dialyzed FCS and glutamine (Silantes, #282946423) for at least two weeks. Then ‘light’ labeled cells were infected with *Ctr* for 24 h and ‘heavy’ labeled cells were left uninfected. Aliquots were treated with staurosporine in the presence of caspase inhibitor (10 µM). Whole cell lysates were obtained by lysing the cells in 1% CHAPS. Precleared lysates were immunoprecipitated for 120 min using either Bak(aa23-38) or Bak(Ab-1) and protein G agarose beads. Before elution, beads from *Ctr*-infected-versus uninfected-sample for the same experimental condition and same antibody were mixed. Bound fractions (eluates) were obtained by boiling the beads at 95 °C in Laemmli-buffer and were run on SDS-PAGE and processed for mass spectrometry. LC-MS/MS analysis of the in-gel digested, co-immunoprecipitated proteins was performed as described previously [37]. Data were analyzed by MaxQuant v 1.5.2.8 using a combined database, consisting of Chlamydia trachomatis serovar and human protein sequences. The false discovery rate at the peptide and protein level was 1 %. Proteomic data are available via PRIDE/ProteomeXchange with identifier PXD011848 (reviewer account details – to be deleted upon eventual publication: username; password: VB0DOcEG).

### SDS-PAGE and Western blotting

Cells were lysed and boiled in Laemmli buffer (Thermo Fisher, #89900). Samples were heated for 5 min. Antibodies used were: Bak(NT), Bak(aa23-38), Bak(Ab-1), Bak(7D10) as above. The following antibodies were purchased from Cell Signaling unless indicated otherwise (targets): GAPDH (Millipore, #MAB374), BAX (#2772), VDAC (#4661), Hsp60 (#4870), SMAC (#2954), Hsp60(*Ctr*) (Enzo Life Sciences, #ALX-804-072), OmpA (Biozol Diagnostica, #LS-C79219), FLAG (Sigma, #F1804), GFP (Roche, #11814460001), BiP (Cell signalling, #3177), Golgin-84 (SantaCruz, #sc-365337), Lamin B (Abcam, #ab45848), Rab7 (Abcam, #ab50533), α-tubulin (Sigma, #T9026). Peroxidase-conjugated secondary antibodies were goat anti-rabbit IgG (Sigma, #A6667), goat anti-rabbit Fc (Sigma, #AP156P), goat anti-mouse IgG (Dianova, #115035166), goat anti-mouse Fc (Sigma, #AP127P), goat anti-rat IgG (Dianova, #112035062) and mouse anti-goat (Dianova, #205035108).

### Single-Molecule Localisation Microscopy (SMLM)

For OmpA staining, cells were seeded on glass coverslips (Marienfeld, 24mm, #1.5-H-117540) in a 6 well-plate and infected with Ctr (MOI=5). Cells were fixed 24 h post-infection for 15 min in 4% PFA, were permeabilized for 10 min in 0.2% Triton X-100 (Sigma) in PBS, and incubated for an additional 30 min in 5% BSA in PBS. Polyclonal goat anti-OmpA (Biozol Diagnostica, #LS-C79219) was used to detect OmpA, followed by AF647-conjugated donkey anti-goat (Thermo Fisher, #A21447). To investigate mitochondrial localization of OmpA, mitochondrial protein Tom20 was co-stained with OmpA using polyclonal rabbit anti-Tom20 (Santacruz, #sc11415), followed by CF680-conjugated donkey anti-rabbit (Biotium, #20418). DNA was stained with Hoechst 33342 (1 mg/ml; Sigma).

### Microscope setup and imaging for SMLM

SMLM data were acquired on a custom built widefield setup described previously. Briefly, the free output of a commercial laser box (LightHub, Omicron-Laserage Laserprodukte) equipped with Luxx 405, 488 and 638 and Cobolt 561 lasers and an additional 640 nm booster laser (iBeam Smart, Toptica) were collimated and focused onto a speckle reducer (Optotune, Dietikon, #LSR-3005-17S-VIS) before being coupled into a multi-mode fiber (Thorlabs, #M105L02S-A). The output of the fiber was magnified by an achromatic lens and focused into the sample to homogeneously illuminate an area of about 1,000 μm2. Alternatively, a single-mode fiber (Omicron, LightHUB) could be plugged into the output of the laserbox to allow TIRF imaging. The laser is guided through a laser cleanup filter (390/482/563/640 HC Quad, AHF) to remove fluorescence generated by the fiber. For ratiometric dual-color imaging of AF647 and CF680, the emitted fluorescence was collected through a high numerical aperture (NA) oil immersion objective (Leica, #HCX PL APO 160×/1.43 NA), split by a 665LP beamsplitter (Chroma, #ET665lp), filtered by a 685/70 (Chroma, #ET685/70m) bandpass filter (transmitted light) or a 676/37 (Semrock, #FF01-676/37-25) bandpass filter (reflected light) and imaged side by side on the EMCCD camera. The color of the individual blinks was assigned by calculating the ratio of the intensities in the two channels. Astigmatism was introduced by a cylindrical lens (f = 1,000 mm; Thorlabs, #LJ1516L1-A) to determine the z position of fluorophores. The z focus was stabilized by an infrared laser that was totally internally reflected off the coverslip onto a quadrant photodiode, which was coupled into closed-loop feedback with the piezo objective positioner (Physik Instrumente). Laser control, focus stabilization and movement of filters were performed using a field-programmable gate array (Mojo, Embedded Micro). The pulse length of the 405 nm (laser intensity 27.5 W cm−2) laser is controlled by a feedback algorithm to sustain a predefined number of localizations per frame. Coverslips containing prepared samples were placed into a custom build sample holder and 500 μl of blinking buffer (50 mM Tris/HCl pH 8, 10 mM NaCl, 10% (w/v) d-glucose, 500 μg ml–1 glucose oxidase, 40 μg ml–1 glucose catalase, 35 mM MEA) was added. To avoid a pH drift caused by accumulation of glucuronic acid in GLOX-buffers, the buffer solution was exchanged after about 2 h of imaging. Samples were imaged until close to all fluorophores were bleached and no further localizations were detected under continuous ultraviolet irradiation.

### Data analysis for SMLM

All data analysis was conducted with SMAP, a custom software written in MATLAB that is available as open source (github.com/jries/SMAP). Installation instructions are found in the README.md, and step-by-step guides on how to use the software to perform all analyses used in this manuscript are available via the Help menu.

### Fitting for SMLM

Astigmatic 3D data were fitted and analyzed as described previously (65). First, z stacks with known displacement of several (15–20) fields of view of TetraSpeck beads on a coverslip were acquired to generate a model of the experimental point spread function. This model was then used to determine the z position of the individual localizations. Free fitting parameters: x, y, z, photons per localization, background per pixel.

### Post-processing for SMLM

The x, y, and z positions were corrected for residual drift by a custom algorithm based on redundant cross-correlation. Localizations persistent over consecutive frames (detected within 35 nm from one another and with a maximum gap of one dark frame) were merged into one localization by calculating the weighted average of x, y and z positions and the sums of photons per localization and background. Localizations were filtered by the localization precision (0–20 nm) to exclude dim localizations. Additionally, poorly fitted localizations were excluded if their log-likelihood (LL) was smaller than the mean (LL)—3 × STD (LL).

Superresolution images were constructed with every localization rendered as a two-dimensional elliptical Gaussian with a width proportional to the localization precision (factor 0.4). The reported mean photons per localization were calculated based on these merged and filtered localizations.

## Acknowledgement

This study was supported by the Deutsche Forschungsgemeinschaft (grant HA 2128 to G.H.). The Core Facility for Electron Microscopy (EMcore) at the University Freiburg Medical Center—IMITATE is registered with the DFG (German Research Foundation) under the reference number RI_00555.

We are grateful to Severine Kayser for excellent technical assistance with TEM.

---

**Fig. S1** Comparisons of predicted OmpA of Chlamydia-species and Simkania negevensis

**A**, Clustal Omega sequence alignment comparisons of various chlamydia species comparing to *Simkania negevensis* and several *Rhabdochlamydia* species which show high level of similarity to *Simkania* OmpA. Red bar indicates the signal peptide, Black arrows indicate the transmembrane beta barrel forming beta strands of Chlamydia OmpA as predicted by Alphafold seen in B. Colour is based on Clustal colouring. Protein identity and amino acid position are indicated. Note the high primary sequence identity among the different OmpA-homologous proteins. **B**, Alphafold predictions of Chalmydia OmpA (Left) and Simkania negevensis OmpA (right). Colouring is from alpha fold based on confidence of prediction (shown with colour key).

**Fig. S2** Lack of PLA-signal in Ctr*-infected BAK-deficient HeLa cells*

BAK-deficient HeLa cells were seeded on cover slips and infected with *Ctr*. 24 h post-infection, cells were treated with ABT-737 (1 μM) and S63845 (500 nM) for 4 h in the presence of the caspase inhibitor QVD-OPh (10 μM). Cells were fixed, permeabilized and processed for PLA using antibodies against BAK (Ab-1(TC-100)) and OmpA. Mitochondria were labeled using antibodies directed against Tom20 (green), and DNA was stained with Hoechst dye (blue). Cells were imaged by confocal microscopy. Data are representative of three independent experiments. Scale bar, 5 μm.

**Fig. S3** BAK interacts with chlamydial OmpA

**A,** experimental design. HeLa cells were differentially labelled in SILAC media (‘light’ or ‘heavy’ amino acids, marked L or H). ‘Light’ cells were infected with Ctr (MOI=5) for 24 h. Two aliquots of cells were additionally treated with staurosporine as indicated and lysed. Two sets of two lysates were combined as indicated. Lysates were precipitated using two different anti-BAK antibodies (both against active BAK, four IP-reactions in total), and four samples were collected. IP-products were analyzed by mass spectrometry. For analysis we focused on proteins identified in all reactions. **B,** OmpA Ibaq values correlate with BAK Ibaq values. Ibaq values of OmpA and BAK obtained by the proteomic analysis (a measure of protein abundance) are plotted against each other. Note the correlation of the values from the two experiments each where the same antibodies were used (#1 and #3, #2 and #4). **C,** Interaction between BAK and OmpA as detected by co-immunoprecipitaiton. *Ctr*-infected HeLa cells (MOI=5) were treated with the combination of ABT-737 (1 μM) and S63845 (500 nM) for 4 h. Mitochondria were isolated, lysed in buffer containing 1% CHAPS, and lysates were subjected to immunoprecipitation using BAK (aa23-38) antibody or an IgG isotype control antibody. Interaction of BAK and OmpA was visualized by immunoblotting for Bak and OmpA proteins.

**Fig. S4** HeLa cell 3D SMLM reveals interactions between OmpA und mitochondria with nanometer-scale resolution

**A,** schematic representation of the analysis. For every OmpA localization we counted the number of Tom20 localizations that are closer than 50 nm in the lateral and 150 nm in the axial direction. OmpA localizations with less than 5 Tom20 neighbors are considered to be part of the background and OmpA localizations with more than 5 Tom20 neighbors are considered to be associated to mitochondria. **B,** OmpA is associated to mitochondria. We normalized the number of OmpA localizations at mitochondria to the number of Tom20 localizations and the OmpA localizations at the background to the area of the image to be independent of the image size. As a control, we shifted and mirrored the OmpA image and calculated the number of neighbors for this randomized negative control. We find that OmpA is associated to mitochondria, and that the number of OmpA molecules for the infected or overexpressed case is substantially larger than the background staining in the wildtype. WT (HeLa cells wt); infected (Ctr-infected HeLa cells for 24 h, (MOI=5)); expressed (48 h OmpA expression in HeLa cells carrying a tetracycline-inducible OmpA). The data for this figure were created with the OmpA_mito_cc.m script that is included in SMAP (https://github.com/jries/SMAP).

**Fig S5** Nanoparticle Tracking Analysis of OMVs

Chlamydial OMVs (upper panel) were labelled with primary anti-OmpA and secondary AF-488 antibodies. The fluorescence was measured with ZetaView with the laser wavelength 488 nm, temperature 24.01 °C. The output gave an OMVs size peak at 118.7 nm and an absolute number of 2.4E+9 particles/ml. Control OMVs (lower panel) were also measured with with ZetaView with the laser wavelength 488 nm, temperature 23.94 °C. The output gave a particle size of 78.4 nm and an absolute number of 9.0E+8 particles/ml.

**Fig S6** OmpA sequence conservation and AlphaFold structure of OmpA

AlphaFold software from DeepMind was used to predict following structures and interactions between OmpA:VDAC2 and OmpA:BAK. **A**, Shown are comparisons of VDAC2 with Bak in a 1:1 stoichiometry. The predicted orientation of the structures are indicated. **B**, Shown are comparisons of VDAC2 and BAK in a 1:1 stoichiometry OmpA and BAK with a 3:3 stoichiometry. The predicted orientation of the structures are indicated

## Notes

### Competing Interest Statement

The authors have declared no competing interest.

